# Characterizing amblyopic perception under naturalistic viewing conditions

**DOI:** 10.1101/2022.10.10.511635

**Authors:** Kimberly Meier, Kristina Tarczy-Hornoch, Geoffrey M. Boynton, Ione Fine

## Abstract

Current assessments of interocular interactions in amblyopia use rivalrous stimuli, with conflicting stimuli in each eye, which does not reflect vision under typical circumstances. Here we measure interocular interactions in observers with amblyopia, strabismus with equal vision, and controls using a non-rivalrous stimulus. Observers used a joystick to continuously report perceived contrast of dichoptic grating stimuli, identical except that the stimulus was contrast-modulated independently in each eye over time. Consistent with previous studies, a model predicting the time-course of perceived contrast found increased amblyopic eye attenuation, and reduced contrast normalization of the fellow eye by the amblyopic eye, in amblyopic participants compared to controls. However, these suppressive interocular effects were weaker than those found in previous studies, suggesting that rivalrous stimuli may overestimate the effects of amblyopia on interocular interactions during naturalistic viewing conditions.

## 1 INTRODUCTION

Amblyopia is a visual disorder clinically characterized as poor visual acuity in one eye even with best optical correction in place. Amblyopia, which affects 1-2% of the population^1^, arises when a child experiences abnormal vision in one eye for a prolonged period of development, typically because of anisometropia (unequal refractive error in the two eyes), strabismus (an eye misalignment), or a combination of the two, resulting in subnormal best-corrected vision^2^. Amblyopia is a consequence of the abnormal progression of cortical development in response to this atypical visual input during key periods of maturation^3^, rather than being directly attributable to optical or physical characteristics of the eye.

Amblyopia results in disruption of a variety of visual functions routinely assessed in the clinic, such as acuity, contrast sensitivity, fusion, and stereopsis (reviewed in ^4–6^). Generally, the mechanisms underlying binocular amblyopic deficits are thought to include *attenuation* of the amblyopic eye signal and abnormal interocular *suppression*, or inhibition, of the signal of one eye by the other, often modelled as some form of gain control^7,8^, e.g. contrast normalization^9–11^.

A wide variety of dichoptic stimuli^12^ have been used to characterize the effects of amblyopia on binocular vision, including gratings and plaids^8,13–15^, moving dots^16^, and letters^17^. However, these tasks have all relied on rivalrous stimuli, presenting dichoptic images that differ in their spatial content, orientation, signal/noise properties, phase etc. across the two eyes.

One concern with this approach is that rivalrous conditions do not reflect the naturalistic viewing conditions experienced in daily life. When both eyes are open (assuming adequate correction of etiological factors, such as proper spectacle prescription and surgical alignment), the visual cortex receives consistent input from each eye over most of the visual scene, as shown in Figure 1A. In regions where the visual scene is consistent across the two eyes, the statistically optimal strategy is presumably to combine signals from the two eyes with a weighting that reflects the relative ‘reliability’ of each eye. However, there are circumstances (e.g. differential occlusion in each eye of a distant object by a nearer object), Figure 1B, where the input between the two eyes differs substantially over a local region of the scene, and any weighted average of the two eyes will be perceptually nonsensical, Figure 1C. In these rivalrous regions, the best strategy is presumably a ‘winner-take-all’ in which the input from the most ‘reliable’ eye dominates. Thus, the mechanisms of attenuation and interocular suppression that operate under conditions of rivalry – as used for most measures of interocular suppression in amblyopia – might easily be very different from those that underlie neural responses under more typical non-rivalrous viewing conditions.

**Figure 1.**
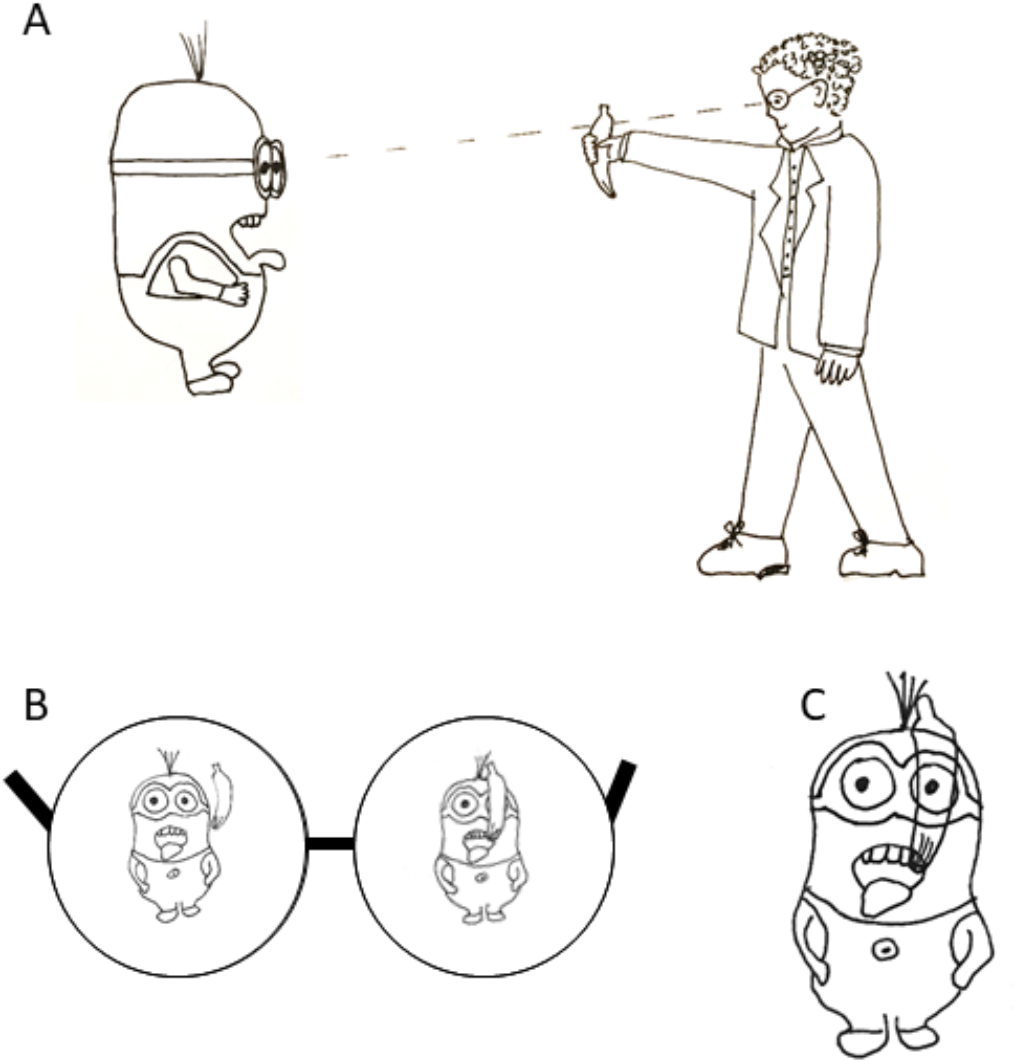
In typical binocular vision (Panel A) most of the visual scene is consistent between the left and right eye (Panel B, the region of Dave the Minion’s overalls), and the statistically optimal strategy is to combine signals from the two eyes with a weighting that reflects the relative ‘efficiency’ of each eye. However in certain circumstances (e.g. partial visual occlusion by a banana) the input received by the two eyes will differ substantially over a local region of the scene (Dave’s goggles) and any weighted average of the two eyes will be perceptually nonsensical (Panel C). In these regions of the scene the best strategy is a ‘winner-take-all’ where one eye’s input dominates.

To address this, we developed a non-rivalrous task for assessing the neural mechanisms underlying amblyopia. Participants were shown dichoptically-presented grating stimuli whose spatial content was identical, except for their contrast, which was modulated independently in each eye over time. Participants dynamically adjusted the position of a joystick lever over time to match their perception of stimulus contrast.

To maximize our ability to examine interocular interactions, we used 2 cpd grating stimuli. This is roughly the peak of the contrast sensitivity function for amblyopic eyes^18^; indeed, amblyopic observers often do not show interocular differences in monocular contrast detection thresholds at this spatial frequency^19^. Thus our choice of a relatively low spatial frequency stimulus was designed to ensure that our measurements primarily reflected suprathreshold interocular interactions^20^ rather than monocular attenuation from a degraded input signal.

We found that, for our non-rivalrous stimulus, a simple model based on *monocular attenuation in the context of binocular stimulation* (k_AE_), and *interocular contrast normalization* (μ_AE_: normalization of the amblyopic by the fellow eye; μ_FE_: normalization of the fellow by the amblyopic eye) could successfully predict response time-courses. Fitted model parameters predicted performance on independent clinical measures of stereoacuity and interocular balance point.

Our estimates of interocular contrast normalization, as measured in our naturalistic task, were considerably weaker than those previously reported for rivalrous conditions, suggesting that the ‘perceptual burden’ of amblyopia under naturalistic viewing conditions may be weaker than has previously been supposed.

## 2 RESULTS

### 2.1 Dynamic contrast estimation task

The stimulus, shown in Figure 2A, consisted of a Gaussian-windowed 2 cpd grating on a mean-luminance grey background. The grating rotated slowly counter-clockwise at 1°/s for the duration of the trial to minimize adaptation. Each trial began with 14 s of *binocular* 1/7 Hz contrast modulation, in which the contrast of the two eyes was identical. Most of the remaining 48 s of the trial was *dichoptic*, such that contrast in one eye modulated at 1/6 Hz while the contrast in the other eye modulated at 1/8 Hz (or vice versa). The periods of these sinusoidal modulations synchronized every 24 s, so each modulation pattern was repeated once during each trial. Embedded in each trial was a short phase of *monoptic* stimulation in which the contrast modulation in one eye ‘dropped out’ and remained at 0% contrast for a single cycle (6 or 8 s), while the other eye continued to modulate. Example contrast time-courses are shown for two trials in Figure 2B.

**Figure 2.**
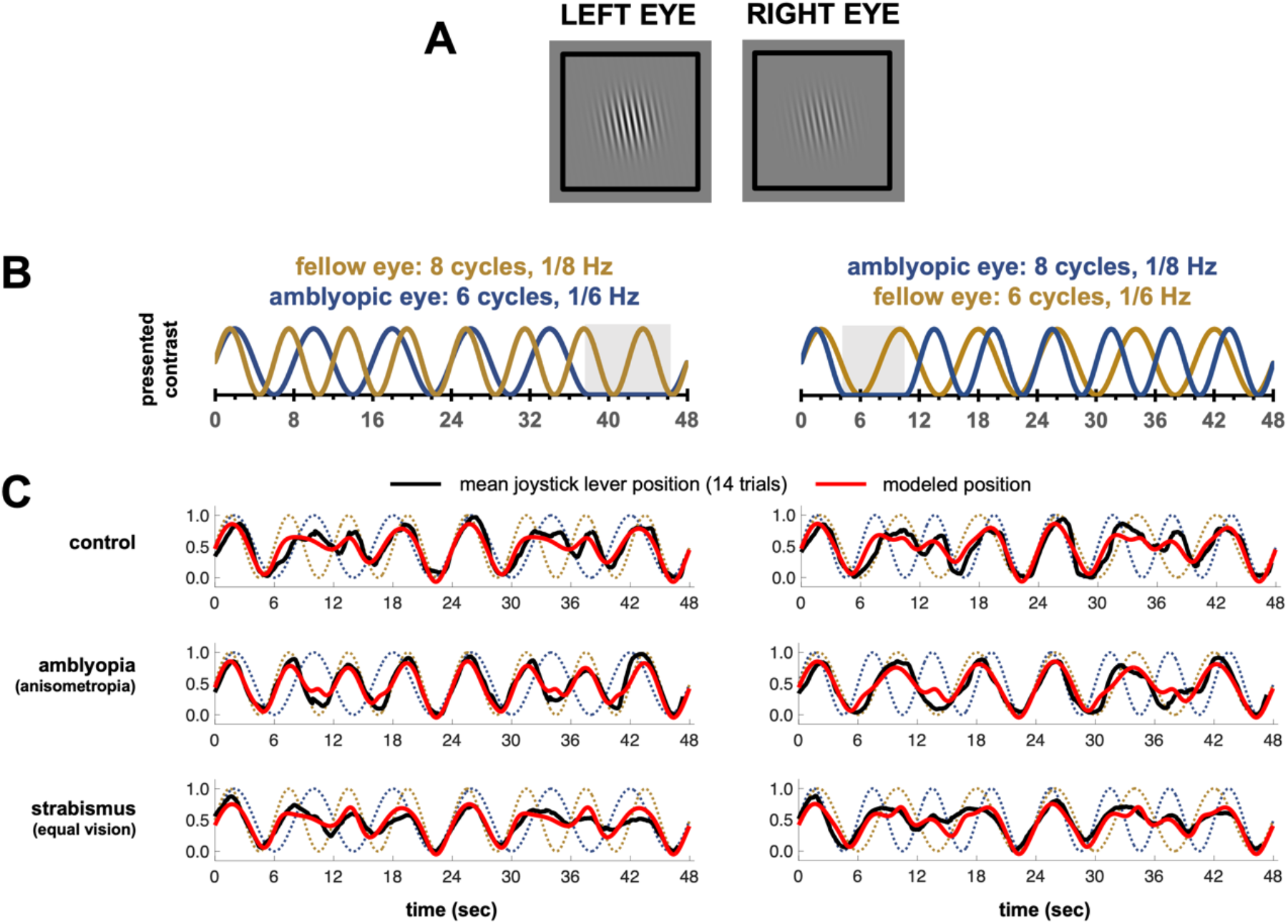
(A) Example dichoptic stimulus. (B) Trial schematic indicating contrast modulation in each eye over time. In half of the trials the fellow eye contrast modulated at 1/8 Hz and amblyopic eye at 1/6 Hz (left panel) or vice versa (right panel). Example monoptic portions embedded in the trial are shown in grey, these regions differed across each trial. (C). Example mean time-courses averaged across all 14 trials per condition (dichoptic periods only), with model fits (based on the full dataset), for typical participants from the control, amblyopic and non-amblyopic strabismus with equal vision groups.

We used a three-stage model fitting procedure to predict joystick position over time as a function of the contrast presented to each eye. The first stage calibrated the relationship between joystick position and perceived contrast using the portion of each trial that was binocular, the second stage modeled monocular attenuation using the portion of each trial that was monoptic, and the third stage modeled binocular interactions using the portions of each trial that were dichoptic.

Figure 2C displays the mean joystick lever position over time averaged over all 14 trials for typical individuals from each group. The dotted lines show the contrasts presented to each eye, the red lines show the model’s predicted joystick position, and the black lines show actual joystick position. Both conditions (contrast modulation 1/8 Hz amblyopic, 1/6 Hz fellow; and 1/6 in amblyopic, 1/8 Hz fellow) are shown. The model accurately predicts perceived contrast over time. Inspection of residuals revealed no obvious systematic deviations between model predictions and joystick position. Example-course data for single trials is shown for the same individual participants in Supplementary Figure 1.

### 2.2 Modeling Stage 1: Joystick vs. contrast calibration

The first modeling stage characterized idiosyncratic deviations between joystick position and presented binocular contrast using a simple transformation between stimulus contrast and joystick position. This consisted of a parameterized static linear function and a short delay between the stimulus contrast and joystick position. Inverting this transformation maps joystick position to contrast, which we call the calibrated joystick position. Best-fitting delay and linear function parameters were found by minimizing the mean square error between the calibrated joystick position and stimulus contrast *C* as a function of time. None of the joystick calibration parameter values, including MSE in calibration fit, differed significantly between groups, see Supplementary Table 1.

### 2.3 Stage 2: Monocular attenuation

Next, data from the monoptic phase of each trial were used to estimate the monocular attenuation weights k_L_ and k_R_, as follows:

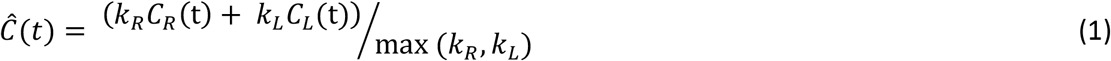

Where *Ĉ*(*t*) is the model prediction of perceived contrast, and *C_R_*(t) and *C_L_*(t) are the contrasts of the stimuli presented to right and left eyes respectively. Because this model was only fit to the monoptic phase of the trials, either *C_R_*(t) or *C_L_*(t) was always 0. The normalization by max (*k_R_*, *k_L_*) meant that either *k_R_* or *k_L_* became 1. We defined the eye with k = 1 as the ‘fellow’ eye, and the eye with k < 1 as the ‘amblyopic’ eye, *k_AE_*, in all observers, with *k_AE_* characterizing linear monocular attenuation in the amblyopic eye. This model is a reduced form of the full model, as described below, with μ_R_ = μ_L_ = 0, and σ = 1.

As shown in Figure 3A and Table 1, *k_AE_* was significantly lower in observers with amblyopia than in controls (*p* < 0.0001), though it is worth noting that there was significant individual variability in participants with amblyopia. In strabismus with equal acuity, values of *k_AE_* were intermediate between amblyopia and controls.

**Figure 3.**
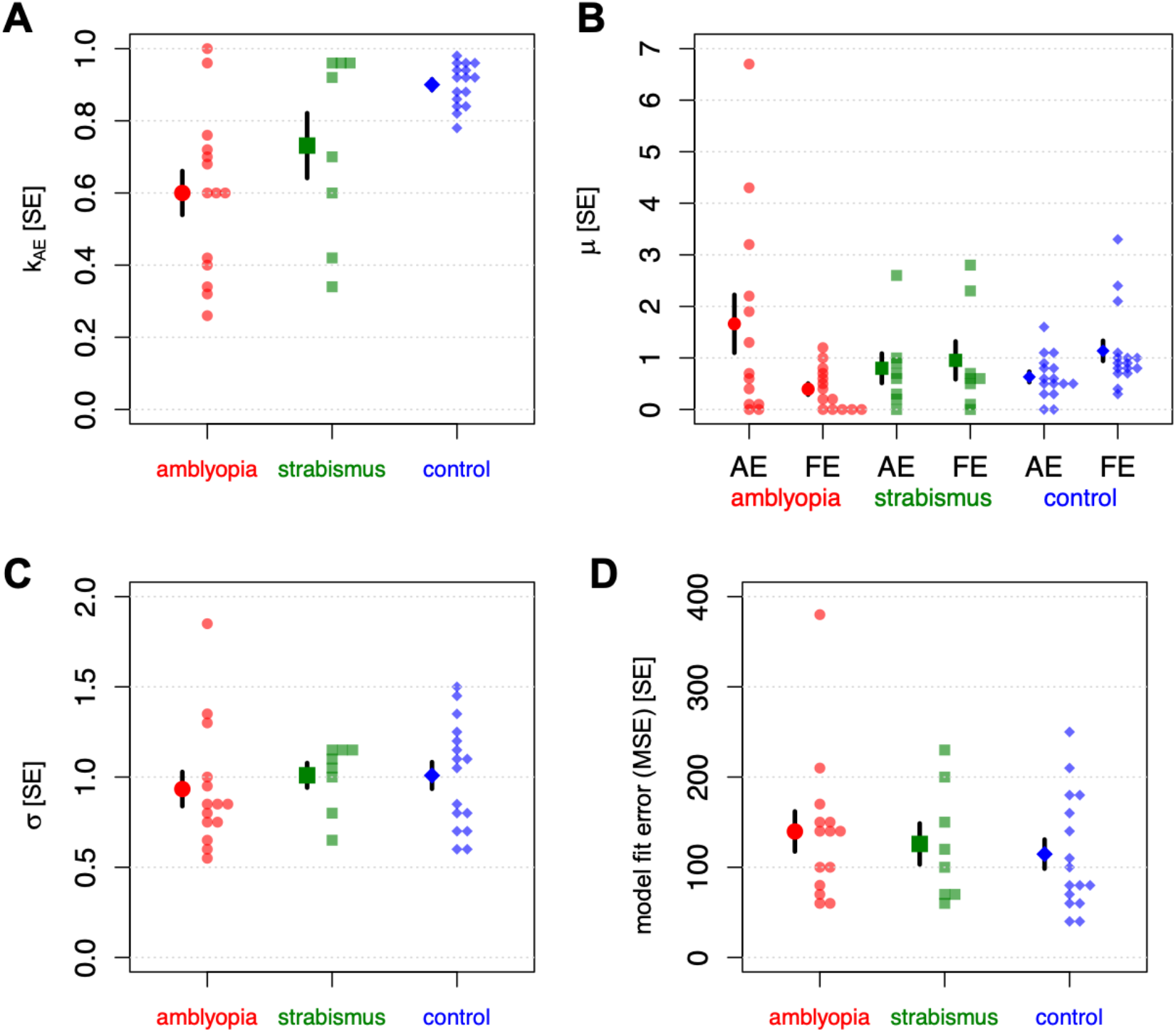
Parameter values used to fit the dynamic contrast estimation task. (A) k_AE_, (B) μ_AE_ and μ_FE_, (C) μ, (D) MSE. Mean values and single standard errors are shown with the larger symbols, individual data points are shown with the smaller symbols. Subjects in the amblyopia group had strabismic, anisometropic or combined amblyopia. Subjects in the strabismus group had equal visual acuity.

**Table 1.**
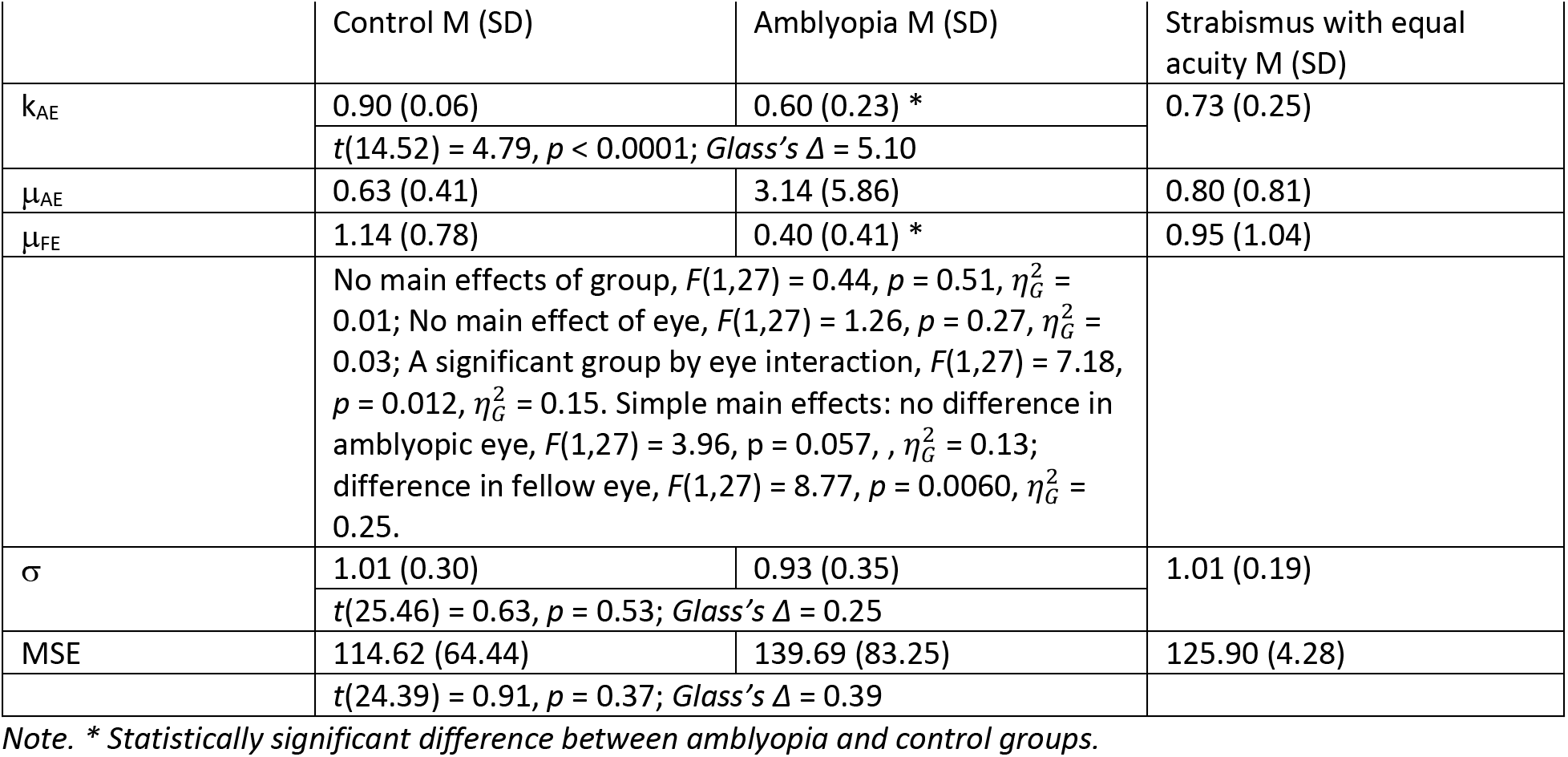
Monocular attenuation and binocular interactions. Mean (SD) values are shown.

### 2.4 Stage 3: Binocular interactions

We then modeled the remaining dichoptic portions of each trial, using a simple model, consisting of attenuation (described by the parameter *k_AE_* estimated in Stage 2), divisive normalization and the linear combination of signals from the two eyes. The parameter *k_AE_* was held fixed, having been estimated from the monocular intervals as described above.

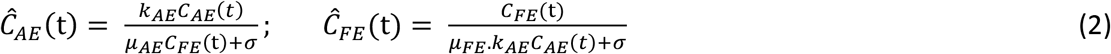

Thus μ_*AE*_ and μ_*FE*_ reflect the extent to which the signal in the designated eye is reduced, or suppressed, by the signal in the other eye; in a ‘perfectly balanced’ system we would expect μ_AE_ = μ_FE_.

Finally, we assume the final perceived contrast is simply the sum of the normalized attenuated values for the left and right eyes (no additional free parameters).

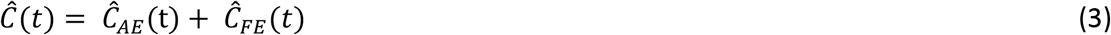

As shown in Figure 3B, and Table 1, there was a significant group by eye interaction (*p* = 0.012), driven by significantly lower μ_FE_ (less suppression of the fellow by the amblyopic eye) in observers with amblyopia compared to controls (*p* = 0.0060): suggesting that the amblyopic eye contributed weakly to contrast normalization in the fellow eye. Interestingly, five participants with amblyopia (36%) have values of zero or near-zero for μ_FE_, a phenomenon not demonstrated to the same extent by participants in the other groups, including those with non-amblyopic strabismus. μ_AE_ (suppression of the amblyopic by the fellow eye) was not statistically different between the groups (*p* = 0.057), though this parameter showed considerable individual variation. Thus in our model the effects of amblyopia on binocular interactions were predominantly described by the amblyopic eye failing to contribute to contrast normalization in the fellow eye, rather than an increase in normalization from the fellow eye to amblyopic eye.

One consideration was that, despite being fit on different data, attenuation and suppression might play a similar role, and trade-off against each other in the model. However the correlation between attenuation (k_*AE*_) and suppression (μ_AE_) across all participants was *r* = 0.13, *t*(37) = 0.80, *p* = 0.43; and within the amblyopia group only, this was *r* = 0.30, *t*(12) = 1.07, *p* = 0.30. Thus, it does not appear that *less* suppression of the amblyopic eye occurs when it is already highly attenuated, as would occur if these parameters traded off against each other in the model.

Finally, neither the saturation constant (σ) or the mean model error (MSE) differed across control and amblyopia groups, Figure 3C and D, and Table 1; observers with non-amblyopic strabismus had similar values.

### 2.5 Assessments of visual function

In addition to the dynamic contrast task described above, we carried out four assessments of visual function, as shown in Figure 4 and Table 2.

**Figure 4.**
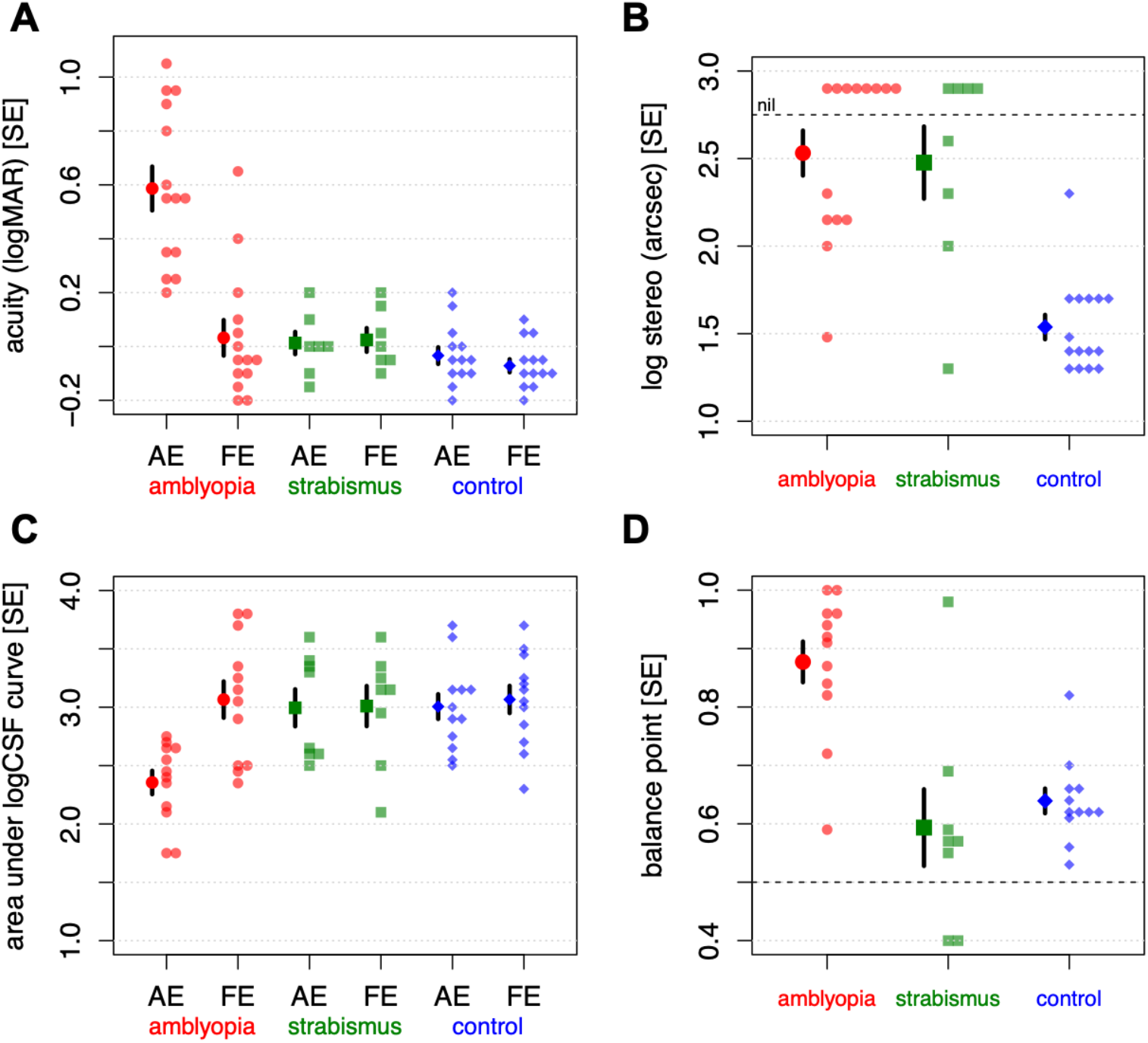
Measures of visual performance. (A) Acuity, (B) Stereo, (C) Area under log CSF curve and (D) Interocular balance point. Mean values and single standard errors are shown with the larger symbols, individual data points are shown with the smaller symbols. Subjects in the amblyopia group had strabismic, anisometropic or combined amblyopia. Subjects in the strabismus group had equal visual acuity.

**Table 2.**
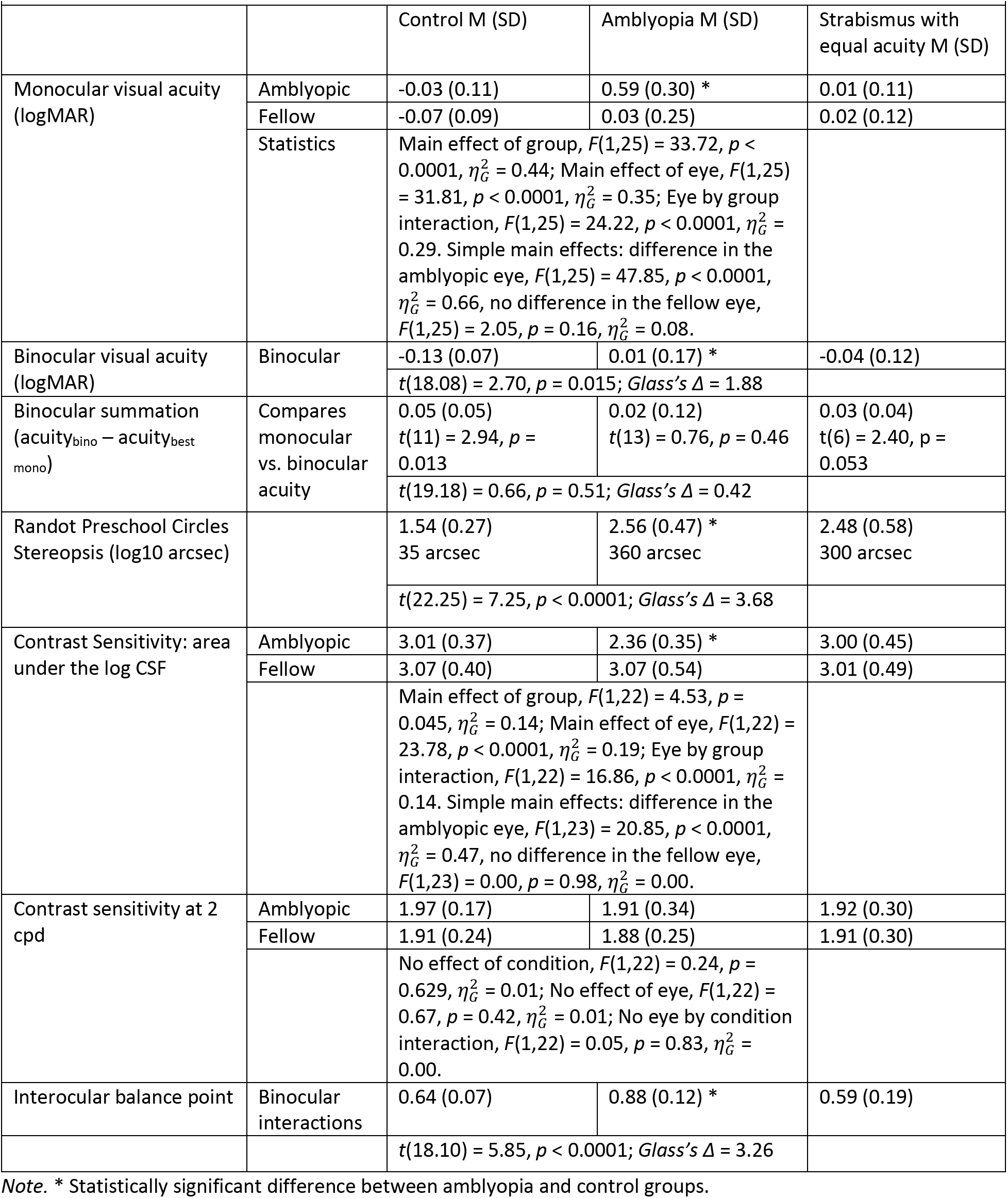
Results for Assessments of Visual Function

#### 2.5.1 Monocular and Binocular Acuity

Amblyopia participants had significantly worse monocular acuity than controls in the amblyopic (*p* < 0.0001), but not the fellow eye (*p* = 0.16), Figure 4A. Although binocular visual acuity was within normal range for observers with amblyopia, measured values were significantly lower than those of control participants (*p* = 0.015). Acuity in observers with non-amblyopic strabismus resembled controls.

Binocular summation (or inhibition) of acuity is defined as the increase (or decrease) in acuity when using both eyes relative to the best monocular visual acuity. Control observers showed binocular summation significantly different from zero (*p* = 0.031), while participants with amblyopia and those with strabismus with equal vision showed no significant change in acuity (in either direction) with binocular viewing (*p* = 0.50 and 0.053, respectively). However, the degree of summation in the control group did not differ significantly from that in the amblyopia group (*p* = 0.51).

#### 2.5.2 Stereopsis

Stereoacuity on the Randot Preschool Circles test is shown in Figure 4B; stereoacuity was significantly worse in observers with amblyopia than controls (*p* < 0.0001). Stereoacuity in strabismus with equal vision resembled the amblyopia group.

#### 2.5.3 Contrast sensitivity

There were no differences in sensitivity at 2 cpd (group: *p* = 0.24; eye: *p* = 0.42; interaction: *p* = 0.83), confirming no group differences in contrast sensitivity at the spatial frequency of our stimulus.

Area under the log contrast sensitivity curve (AUC) is shown in Figure 4C. Reflecting acuity losses, participants with amblyopia had a significantly smaller AUC in the amblyopic eye (*p* < 0.0001) but not the fellow eye (*p* = 0.98) compared to controls, due to lower sensitivity for higher spatial frequencies in the amblyopic eye. Strabismus with equal acuity showed sensitivity on par with controls.

#### 2.5.4 Interocular balance point

The interocular balance point was assessed by finding the interocular contrast ratio that a participant requires to report seeing the letters in the left and right eye with equal probability {Birch et al., 2016, #44658; Kwon et al., 2015, #183369}. A typical control observer will have a balance point of about 0.5, indicating an equal amount of contrast is necessary in each eye, and a typical observer with amblyopia will have a balance point greater than 0.5, for example, a balance point of 0.7 indicates 70% contrast in the amblyopic eye is necessary for an observer to be equally likely to report the letters shown to the amblyopic and fellow eye.

Amblyopic observers had balance points that were significantly higher than those of control observers (*p* < 0.0001), Figure 4D. Within participants with amblyopia, there was no significant relationship between balance point and acuity, *r* = 0.12, *t*(10) = 0.38, *p* = 0.72, indicating these balance point values are not simply a function of reduced acuity in the amblyopic eye. All but one participant with strabismus and equal acuity had balance point values within the control range.

#### 2.5.5 Summary

In summary, our clinical determination of control, amblyopia and strabismus with equal acuity were reflected in functional tests of visual performance. Participants with amblyopia showed reduced acuity in the amblyopic eye (unsurprisingly, given that they were clinically identified based on acuity), lower contrast sensitivity, shifted interocular balance and reduced stereopsis. Participants classified as non-amblyopic strabismus with equal acuity showed unimpaired acuity (by definition), contrast sensitivity and interocular balance, but did show reduced stereopsis.

### 2.6 The relationship between model parameters and assessment of visual function

Using linear regression, with either Group *or* k_AE_ as predictors, we found that both Group and k_AE_ were significant predictors for all four assessments of visual function: amblyopic eye acuity, stereoacuity, amblyopic eye contrast sensitivity, and interocular balance point (first two columns in Supplementary Table 2). Next we used a nested model F-test to determine whether k_AE_ significantly improved predictions *after* Group had been included as a categorical factor. k_AE_ improved predictions for the two measures of binocular interactions: stereoacuity and interocular balance point. Thus, even after accounting for group membership, k_AE_ was still significantly correlated with stereoacuity and interocular balance point, Figure 5, such that lower k_AE_ values predicted poorer stereoacuity and a more asymmetric balance point within groups.

**Figure 5.**
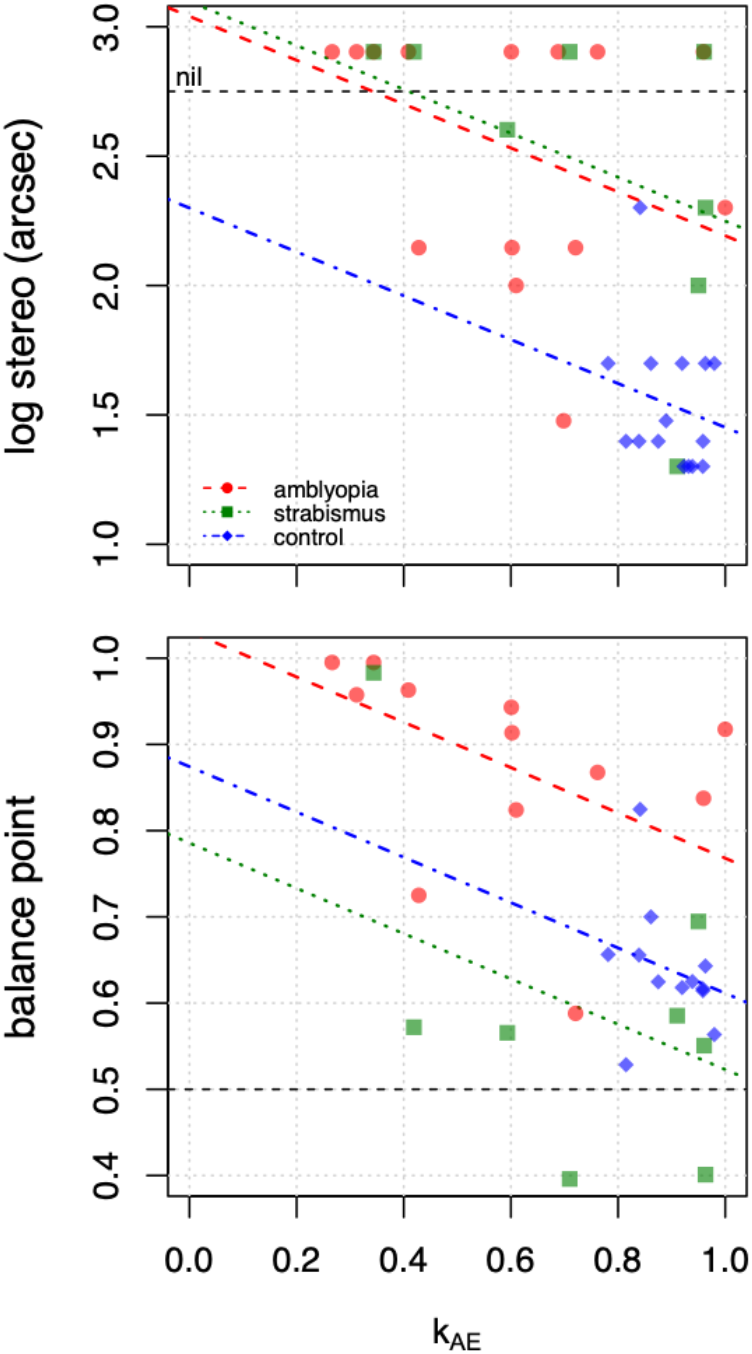
The relationship between k_AE_ and stereoacuity (top) and balance point (bottom) for the regression models described in-text.

The same analysis was performed using our estimated parameters for binocular imbalance, μ_AE_ and μ_FE_ (Supplementary Table 3). While μ_AE_ showed significant relationships with acuity and contrast sensitivity, and μ_FE_ with acuity and interocular contrast ratio, neither of these parameters significantly predicted performance for any of our visual assessments after controlling for group (Supplementary Table 3).

### 2.7 Reduction of task length

Across participants, joystick lever position *J* was well-correlated during the first and second half of each individual trial (dichoptic trial portion only) within all observers (median *r* = 0.66, SD = 0.12, range = 0.29 to 0.87), indicating that most participants tracked perceived contrast over time with reasonable consistency, see Supplementary Figure 1.

We were interested in whether we could obtain reliable parameter estimates for shorter trials – something that would be useful in a clinical environment. We re-fit our models using a reduced dataset consisting of 24 trials, each 38 seconds in length. The correspondence between the original estimates from the full dataset and the estimates for this reduced dataset were extremely high, are shown in Supplementary Figure 2. There was a near-perfect relationship for k_AE_ (r = 0.97), μ_AE_ (r = 0.98), and σ (r = 0.96). The correlation was smaller, but still high, for μ_FE_ (r = 0.75). All results of statistical tests between amblyopia and controls led to identical conclusions. Thus, reasonable parameter estimates can be obtained in approximately 15 minutes of data collection.

### 2.8 Simulating balance point and masking effects

Using the normalization model of equations 1–3 and the mean best-fitting parameters for each group, we simulated our model’s response across a variety of tasks/stimuli previously examined in the literature. These simulations, based on parameter values estimated under non-rivalrous conditions, consistently *under* predicted the effects of amblyopia for experimental data in rivalrous tasks, consistent with the notion that interocular interactions are larger for rivalrous stimuli.

We began by simulating the dichoptic letter chart balance point^17,21^ collected in our observers. Our simulations predicted a balance point of 0.68 for our amblyopic observers, lower than our experimentally measured balance point of 0.88 (control and non-amblyopic strabismus observers had simulated balance points of 0.49 and 0.57 respectively, compared to obtained values of 0.64 and 0.59). Thus, within the same amblyopic observers, we observed greater interocular suppression for rivalrous letters than for the non-rivalrous gratings used in our main paradigm.

Next, we simulated predictions for the cyclopean phase combination balance point^8^ – the perceived phase of a grating produced by presenting opposite phase-shifted sine waves in each eye. This task might be considered ‘semi-rivalrous’ – the input in each eye is different, but tends to be integrated as a weighted average. For control observers, our simulations predicted that roughly equal contrast would be needed in each eye to produce a phase shift consistent with equal perceived contrast in each eye (equivalent to the balance point), a finding that matched measured values of Ding et al.^8^. For observers with amblyopia, at the very lowest contrasts, our simulation results matched that of Ding et al.: roughly twice the contrast was needed in the amblyopic eye to reach the balance point. However at higher contrasts, our model predicted a balance point when the amblyopic eye’s contrast was 2.4x that of the control, while Ding et al.’s experimental data found that a 5-40x increase in contrast was needed to reach balance. Thus, the measured effects of amblyopia in their rivalrous grating balance task were far larger than those predicted by our model.

Finally, we modeled the elevation in contrast required to see a grating in the amblyopic eye when masked by bandpass noise in the fellow eye^22,23^. For controls, our simulations produced dB threshold elevations of about 1.7 across the two experiments, far lower than the observed range of 8 to 13.7 with rivalrous noise stimuli. For observers with amblyopia, our simulations predicted threshold elevations of about 7.6, whereas published experimental values fall between 13.4 and 17^22–24^. Once again, interocular interactions seem to be larger for rivalrous stimuli than predicted by our model, using parameters estimated using non-rivalrous stimuli.

## 3 DISCUSSION

Here we describe an intuitive, efficient, non-rivalrous task for assessing interocular interactions in which observers continuously report the perceived contrast of dichoptic grating stimuli that are identical, except that stimuli are contrast-modulated independently over time in each eye. Participants were observers with amblyopia, strabismus with equal vision, and controls. We fit a model to observers’ responses that agnostically identified the amblyopic eye and allowed us to estimate attenuation of the amblyopic eye and divisive normalization across the two eyes. We found that observers with amblyopia showed greater attenuation and reduced normalization of the fellow by the amblyopic eye compared with controls. Normalization of the amblyopic eye by the fellow eye did not differ significantly between groups.

We found significant monocular signal attenuation (k_AE_) of the amblyopic eye with regard to contrast perception. Although this is consistent with previous findings^20^, it is somewhat surprising for two reasons. First, we found no monocular deficits in the qCSF estimates of sensitivity at 2 cpd, the spatial frequency of our stimulus. Second, our stimulus contrast was primarily suprathreshold, and monocular contrast perception above threshold in the amblyopic eye is generally near-normal for contrast matching and contrast estimation tasks^25,26^.

One possible explanation for this finding of monocular attenuation for suprathreshold stimuli is that during the trial phases used to estimate monocular attenuation of the amblyopic eye, the fellow eye was open and viewing a mean-luminance screen along with a fusion-lock frame. The presence of a foveal signal (albeit non-overlapping) and/or a mean contrast signal in the fellow eye (as compared to patching one eye) may have resulted in attenuation of the amblyopic eye.

Another possibility is that contrast constancy^27^ may break down in amblyopia when monocular periods of stimulation are embedded in the context of dichoptic stimulation – our periods of monocular stimulation were relatively brief (6-8 s), with no cue to their onset. A loss in suprathreshold sensitivity for contrast-matching under dichoptic conditions has previously been shown^20^, and, similar to our study, could be modeled as a combination of input attenuation and increased suppression. Thus, *monocular* attenuation at a given moment in time may be dependent on the previous *binocular* context. This finding is consistent with a variety of studies showing adaptive response normalization across short and long timescales across a wide variety of domains that includes contrast^28–30^.

We found that individual differences in monocular attenuation were significantly related to all four measures of visual function (acuity, stereopsis, contrast sensitivity, and balance point). However, after controlling for group (amblyopia, strabismic with equal vision, or control), only the two measures reflecting binocular interactions – stereoacuity and balance point – remained significantly correlated with attenuation, such that greater attenuation of the amblyopic eye was associated with worse stereoacuity and a more asymmetric balance point. The relationship between attenuation and these two measures may be directly causal. Balance point reflects an interocular contrast ratio, and so is expected to be directly related to perceived contrast. With respect to stereoacuity, reducing the contrast of the image shown to one eye elevates stereoacuity thresholds in control observers^31^, while reducing the contrast of the image sent to the fellow eye improves stereoacuity thresholds in amblyopic observers^32^, consistent with the idea that monocular contrast attenuation interferes with stereoscopic processing.

We saw a significant reduction in the normalization of the fellow eye by the amblyopic eye in amblyopic observers (μ_FE_). In fact, in 5 of our 14 observers with amblyopia we obtained μ_FE_ values at or near zero, indicating no influence at all of the amblyopic eye on the fellow. This is consistent with converging evidence from a variety of psychophysical results in humans^8,22–24^ and electrophysiological evidence from non-human primates^33^ suggesting that amblyopic suppressive imbalance may not be driven by excessive suppression by the fellow of the amblyopic eye (as was previously assumed, and is the basis of many traditional therapies), but rather may reflect a weakened ability of the amblyopic eye to exert influence over the fellow eye. We did not see significantly different normalization of the amblyopic eye by the fellow eye (μ_AE_), though these estimates varied widely among observers with amblyopia.

Our finding that neither attenuation nor normalization in amblyopes predicted visual acuity (after accounting for group) was somewhat surprising, and implies that visual acuity may not adequately summarize amblyopic function^34^. Our task focused on stimuli at 2 cpd, a spatial frequency where contrast sensitivity is almost unimpaired. It seems plausible that attenuation and/or normalization would vary as a function of spatial frequency, as has been shown for rivalrous stimuli^8,17,22,35^, so these parameter estimates might be more strongly correlated with visual acuity when measured using higher frequency stimuli.

Interocular normalization measures (whether from amblyopic to fellow, μ_FE_, or vice versa, μ_AE_) did not predict performance on any of the visual functions we measured, perhaps because most (apart from stereoacuity) were monocular or rivalrous binocular tasks.

Finally, using our normalization model we simulated responses across a variety of tasks and stimuli. Interocular interactions in rivalrous tasks^8,17,21–24^ are consistently larger than predicted by our model, suggesting that the impact of amblyopia may be exaggerated when using rivalrous stimuli as compared to more naturalistic non-rivalrous stimuli.

Collectively, these results are consistent with the intuition that there is likely to be a fundamental difference between the *contrast normalization* that occurs when the signals between the two eyes are consistent (which likely reflects a form of weighted neural averaging/integration) and the interocular interactions that occur when the signals between to the two eyes are locally (as occurs for partial occlusions) or wholly rivalrous, which may result in a form of ‘winner take all’ suppression.

It should be emphasized that the model described here does not attempt to explain the full complexity of interocular interactions. It is likely that our simple model would fail to fit data using the same paradigm but using orthogonal gratings in each eye: consistent with the observation that models of amblyopia using rivalrous stimuli tend to be more elaborate than the model described here^7,8,13,16,36^. However, our model does succinctly describe interocular interactions under naturalistic conditions that favor integration rather than suppression, which may be more reflective of ‘real world’ vision than rivalrous tasks. It should also be noted that our model cannot describe the behavior of an observer with equal vision and alternating strabismus (and therefore alternating suppression) who switches fixation during the task; this would require different parameters for μ during right vs. left eye fixation (e.g. high μ for right and low for left during left eye fixation, and vice-versa).

A key advantage of our contrast-perception paradigm is that it can easily be generalized to a wide variety of stimuli: motion, faces, letters, natural scenes, and so on. Although tasks have been developed to assess and compare the effects of amblyopia across a variety of stimulus domains, it can be difficult to generate pairs of tasks which allow one to be confident that differences in parameter estimates are due to differential effects of amblyopia in brain regions associated with different visual features, rather than differences in the stimulus (e.g. moving dots vs. stationary Gabors) or task (e.g. direction of motion vs. orientation judgments). Our general paradigm can be used to compare parameter estimates across stimuli matched for spatiotemporal frequency content, for example: moving vs. counter-phase flickering gratings, faces vs. scrambled faces, or natural vs. wavelet-scrambled movies^37^. Our paradigm is also easily extendable to the neuroimaging domain (either EEG/MEG or MRI), allowing one to directly compare neural and perceptual time series data.

Another advantage of our paradigm is its efficiency. A multiplicity of dichoptic tasks have been developed to quantify how amblyopia affects the relative strength of eye signals^8,12–17,38^. Many of these have been validated in children^21,39,40^, making them useful research and clinical tools for understanding how interocular suppression changes over time as a function of treatment^41–43^. However these paradigms tend to be either extremely time consuming, require visibility matching^35^, or provide a single measure that cannot successfully isolate both attenuation and suppression (for a brief discussion, see^17^). Our paradigm collects rapid continuous adjustment measurements and is highly efficient: with 15-30 min of data collection, it is possible to reliably fit a model with six-to-seven free parameters that reliably describes both attenuation and interocular suppression. Given that most children today have extensive experience using joysticks and gamepads, it seems likely that this task could be modified to measure the impact of amblyopia therapy in relatively young children.

## 4 METHODS

This study was approved by the University of Washington’s Institutional Review Board, and carried out in accordance with the Code of Ethics of the Declaration of Helsinki. Informed written consent was obtained prior to conducting the experiments.

### 4.1 Participants

Participants were recruited from the greater Seattle, WA community. Participants wore their habitual visual correction during the experiment, if needed.

#### 4.1.1 Amblyopia and/or strabismus

Ophthalmological histories and diagnoses were confirmed by an ophthalmologist (author KTH) and all participants with amblyopia or strabismus underwent an ophthalmological assessment, including retinoscopy and alignment/cover testing. Observers with ≥ 0.2 logMAR interocular acuity difference were classified as having anisometropic amblyopia in the presence of ≥ 1 D spherical or ≥ 1.5 D astigmatic difference in best-corrected refraction, strabismic amblyopia in the presence of heterotropia at near and/or far, and mixed amblyopia if both criteria were met. Observers with < 0.2 logMAR interocular acuity difference in the presence of heterotropia were classified as *non-amblyopic strabismus with equal vision* and are analyzed as a separate group (this group can include both observers with successfully treated amblyopia and observers who had never had amblyopia). In total, 14 participants had amblyopia (*M* age = 50.5 years, *SD* = 19.4, *range* = 18-75); and 8 participants had strabismus with equal vision (*M* age = 40.2 years, *SD* = 18.3, *range* = 18-65). Information about these participants is detailed in Supplementary Table 4. All participants wore their habitual vision correction during testing.

#### 4.1.2 Controls

A total of 16 healthy control participants with no history of vision disorder participated in this experiment (*M* age = 34.6 years, *SD* = 16.4, *range* = 18-69). Because we hoped to capture natural variation in typical development in a group of observers wearing their habitual correction, we did not apply any exclusion criteria to controls based on their performance on any visual assessment tasks. Due to disruptions by the COVID-19 pandemic that occurred when we began running this experiment, 5 control participants do not have data available from all visual assessment tasks. Rather than excluding these participants, we include their available task data, and report when these data are unavailable.

#### 4.1.3 Equipment

Participants viewed stimuli presented on a custom-built mirror stereoscope. A forehead and chin rest were used to stabilize head position. Two mirrors angled at 45° and 135° were positioned in front of the participants’ eyes, each reflecting the image of one of two LED-backlit LCD monitors (Philips 328P6AUBREB) positioned to the left and right of the participant at a viewing distance of 1.36 m. Prior to conducting each task, participants carried out a crosshair alignment task to ensure proper image alignment in the stereoscope. Participants with strabismus who experienced difficulty during the alignment task (n = 3) were manually aligned by the experimenter based on their known deviation; because of the relatively large area and low spatial frequency of the stimulus, extremely precise alignment is not required for this task, so we include their data.

Monitors subtended 28.9 × 16.5 deg and provided the only source of light in the room. Monitors were linearized with minimum and maximum luminance levels of 0.28 cd/m^2^ and 470 cd/m^2^, respectively. Average luminance of the gratings and the mid-grey background was 235 cd/m^2^. This display system was used for contrast sensitivity testing and the main experiment dynamic contrast task.

Custom-built MATLAB scripts (R2021a, The Mathworks, Inc.) using the Psychtoolbox extension version 3.0.15^44–46^ were used for stimulus presentation.

### 4.2 Dynamic contrast task

Each monitor screen showed a 2 cpd grating patch (Gaussian-windowed at 4 deg; full size 13 × 13 deg) on a mean-luminance grey background (Figure 2A). The grating was always presented at an orientation of 90° on the first frame and rotated slowly counter-clockwise at 1°/sec for the duration of the trial to minimize adaptation. Gratings in each eye slowly modulated between 0 to 100% contrast in a sinusoidal fashion at the frequencies described below. A black frame (13 × 13 deg) surrounded the stimulus in each eye, acting as a fusion lock. This frame remained on the screen at full contrast for the entire trial duration as well as during inter-trial rest periods.

Each trial was 62 s long, and consisted of three different phases of stimulation: binocular, dichoptic, and monoptic. Each trial began with 14 s of *binocular* 1/7 Hz contrast modulation, in which the contrast of the two eyes was identical. Most of the remaining 48 s of the trial was *dichoptic*, such that contrast in one eye modulated at 1/6 Hz while the contrast in the other eye modulated at 1/8 Hz. The periods of these sinusoidal modulations synchronized every 24 s, so each modulation pattern was repeated once during each trial (Figure 2B).

During each trial there was a short phase of *monoptic* stimulation (examples highlighted in grey, Figure 2B). During this monoptic phase, the contrast modulation in one eye ‘dropped out’ and the stimulus in that eye remained at 0% contrast for a single cycle, while the other eye continued to modulate. These monoptic phases allowed us to measure monocular contrast response functions in the context of previous dichoptic stimulation.

After a trial was finished, participants were shown a screen with the number of trials they had completed so far. Participants could press a joystick button to initiate the next trial, or verbally ask the research assistant to initiate the next trial. Participants conducted a total of 28 trials in a pseudo-randomized order such that all possible combinations of dichoptic presentation (left eye 1/8 Hz and right eye 1/6 Hz; left eye 1/6 Hz and right eye 1/8 Hz) and timing of the monoptic phase (which 1 of 14 possible cycles was dropped) were presented once.

The participant’s task was to adjust the position of a joystick lever to report the contrast of the stimuli on the screen, such that the lowest position indicated 0% contrast, the highest position indicated 100% contrast, and positions in between corresponded to intermediate contrasts. Participants were encouraged to use the full range of positions on the joystick. Participants were told that the contrast would not always be predictable, and it was stressed that they should not try to anticipate what the contrast would be but should instead focus on reporting what they see in the moment and not to worry if there was a small delay in what they were shown and what they reported. We did not give participants specific instructions on what to do with their eyes except to tell them to look in the middle of the screen. The joystick’s position was sampled at 30 Hz.

Prior to conducting this task, participants were trained on using the joystick lever to report perceived contrast. Training consisted of two phases: (1) moving the joystick lever to control physically-presented binocular contrast on the screen to become familiar with the range of possible contrast values and their corresponding joystick positions, and (2) practice trials of binocularly-presented contrast modulating at 1/7 Hz for 21 s, followed by visual feedback consisting of a graph showing physical contrast vs. joystick position over time. Once participants were comfortable with training and seemed to be reporting physical contrast with reasonable accuracy (a minimum of three training trials), the experiment commenced.

### 4.3 Modeling of the dynamic contrast task

We used a three-stage model fitting procedure to predict joystick position over time as a function of the contrast presented to each eye. The first stage calibrates the relationship between joystick position and perceived contrast, the second stage models monocular attenuation, and the third stage models binocular interactions.

#### 4.3.1 Stage 1: Joystick vs. contrast calibration

Our first modeling stage was designed to characterize the relationship between joystick position and presented binocular contrast.

We used the last 10 seconds of the 14-second binocular trial portion to calibrate the relationship between each observer’s joystick position (J) and binocular contrast over time. This was done by minimizing the mean square error between the calibrated joystick position *Ĵ* and presented contrast *C* as a function of time *(t)*.

Where:

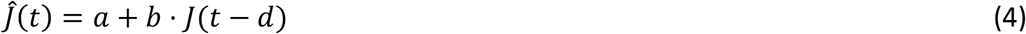

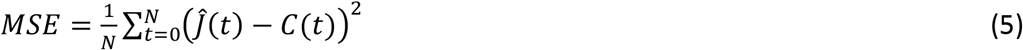

*J* is the joystick lever position (0-1), *d* reflects a time delay between stimulus presentation and observer response, and *a* and *b* represent a linear scaling between joystick location and calibrated joystick position *Ĵ*. A penalization function prevented *d* > 4 seconds, *b* < 0, to avoid behaviorally unrealistic degenerate solutions. Having estimated *d*, *a*, and *b*, these parameters were held fixed for the second two stages of modeling.

#### 4.3.2 Stage 2: Monocular attenuation

Next, data from the monoptic phase of each trial were used to estimate the weights k_L_ and k_R_, as follows:

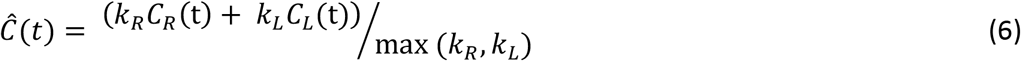

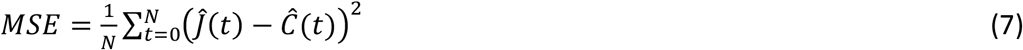

*Ĉ*(*t*) is the model prediction of perceived contrast, *C_R_*(t) and *C_L_*(t) are the contrasts presented to left and right eyes respectively, and *Ĵ(t)* is the calibrated joystick position over time, as described in equations 4 and 5.

Because this model was only fit to the monoptic phase of the trials, either *C_R_*(t) or *C_L_*(t) was always 0. We enforced the constraint that max (*Ĉ*(*t*)) = 1. The presence of max (*k_R_*, *k_L_*) in the denominator, meant that this was achieved by finding the best-fitting solution for which either *k_R_* or *k_L_* was 1, with the other parameter, which always less than 1, representing relative attenuation in the amblyopic eye.

For control and strabismus with equal vision participants, we defined the eye with k = 1 as the ‘fellow’ eye *k_FE_*, and the eye with k < 1 as the ‘amblyopic’ eye *k_AE_*. In all but one case of amblyopia, our modeling successfully identified the amblyopic eye. The participant for whom the amblyopic eye was misidentified had k = 0.93 in the fellow eye; we assigned this participant k = 1 for both eyes at stage 2, and treated the clinically identified eye as amblyopic. Having estimated *k_AE_* and *k_FE_*, these parameters were held fixed for the third stage of modeling.

#### 4.3.3 Stage 3: Binocular interactions

We then modeled the remaining dichoptic portions of each trial, during which contrast differed across the two eyes, using a simple three stage model, consisting of attenuation (parameters held fixed, having been estimated in Stage 2), divisive normalization (such that activity from each eye reduced the gain for the other eye), and a summation of signals from the two eyes.

Linear monocular attenuation was modeled using parameters *k_AE_* and *k_FE_*, using values previously estimated using the monocular portion of each trial, in equations 6 and 7. Since by design *k_FE_* was set to 1:

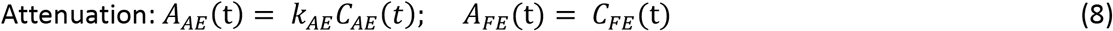

Interocular interactions were modeled using contrast normalization, μ_*AE*_ and μ_*FE*_ to reflect the extent to which the signal in one eye is reduced by the signal in the other eye:

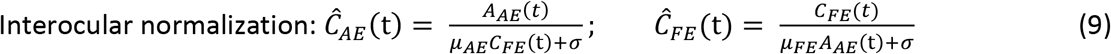

Finally, we assume the final perceived contrast is simply the sum of the outputs of the left and right eyes (no free parameters):

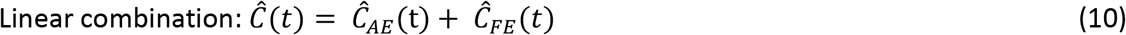

One participant with amblyopia obtained an outlying value of μ_AE_ = 22.3; this value is excluded from the plot and reported statistics.

Thus the parameters *a*, *b*, *d* were estimated based on binocular trial phases, k_AE_, k_FE_, were estimated based on monoptic trial phases, and μ_AE_, μ_FE_, and σ were estimated based on dichoptic trial phases.

### 4.4 Simulating other task predictions

Using the mean group best fitting parameters from our model it is possible to simulate our model’s response to other tasks and stimuli. Using the normalization model above we simulated three tasks/stimuli: (1) our balance point data measured using a dichoptic letter chart (the interocular contrast ratio that a participant requires to report seeing the letters in the left and right eye with equal probability), (2) a balance point as estimated by Ding et al. based on the perceived phase of a 2.72 cpd cyclopean sinewave grating produced by presenting opposite phase-shifted sinewaves to each eye^8^, and (3) the elevation in contrast required to see a grating in one eye with a noise mask in the other eye^22,23^.

To simulate balance points we found the contrast in the amblyopic and fellow eyes that produce equal perceived contrast across both eyes. To simulate the dichoptic letter balance measurements collected in our paper we implemented the experimental constraint that the contrast across the two eyes summed to 1, and used function minimization to find the contrasts that resulted in equal perceived contrast across both eyes. To simulate the cyclopean grating balance measurements, the contrast of the non-dominant eye was fixed to reported experimental values^8^, ranging from 0.0075 to 0.96, and used function minimization to find the contrast of the fellow eye that resulted in the simulation predicting equal perceived contrast across both eyes.

To simulate the effects of masking, we first found the perceived contrast that corresponded to the reported experimental monocular contrast thresholds in the amblyopic (or non-dominant) eye^22,23^. We assumed that a grating in the amblyopic eye would be visible whenever it reached that perceived contrast: and defined this as the *perceived contrast threshold*. We then fixed the contrast in the fellow eye at the reported experimental contrast of the mask and used function minimization to find the contrast in the amblyopic eye that was necessary to reach the *perceived contrast threshold*. Using the metric of these studies, we quantified the elevation in threshold (TE) produced by the mask as:

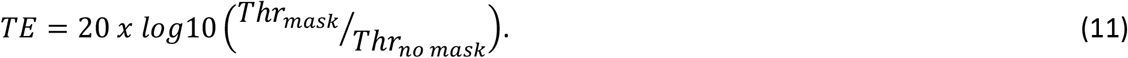

### 4.5 Assessments of visual function

We carried four assessments of visual function, as follows.

#### 4.5.1 Visual acuity

High-contrast visual acuity was assessed binocularly and monocularly while the other eye was occluded using the Acuity letters (5-letter optotype row) program of the Freiburg Visual Acuity Test (FrACT^47,48^) at a 1.5 m viewing distance. Black letters (0.45 cd/m^2^) were displayed on a white background (480 cd/m^2^) using an iMac with a 5K Retina display. The best measurable acuity on this set-up was −0.22 logMAR (20/12 Snellen).

Due to disruptions in data collection, acuity data were not collected from 3 control participants and 1 non-amblyopic strabismus observer who had 20/20 vision in both eyes according to their most recent optometrist documentation.

#### 4.5.2 Stereoacuity

Stereoacuity was assessed using the circles portion of the Randot Stereotest (Stereo Optical Co., Inc.). For analysis purposes, participants with nil stereoacuity are assigned a value of 800 arcsec; all analyses are conducting using log10(arcsec) values. Stereopsis data could not be collected from 1 control participant. Eight of 14 participants with amblyopia and 4 of 8 of participants with non-amblyopic strabismus had no measurable stereopsis using the Randot Circles; these participants are assigned a value of 800 arcsec for analysis purposes.

#### 4.5.3 Contrast sensitivity

Contrast sensitivity (monocular and binocular) was assessed using the Quick CSF method^49^ with 120 trials. Monocular contrast sensitivity was assessed dichoptically, while the other eye was shown a mid-grey background. This task characterizes several useful descriptors of an individual’s contrast sensitivity function; for simplicity, we used log CSF sensitivity at 2 cpd, and the area under the log CSF curve (AUC) as summary measures of contrast sensitivity.

Data were not available from 4 control participants and we could not estimate a CSF in two participants with amblyopia. It was not clear whether they were unable to see the stimulus with either eye, or whether they misunderstood the task.

#### 4.5.4 Interocular balance point

Balance point was assessed using a dichoptic letter chart (described by^17,21^). Briefly, the contrast of dichoptically-presented letters is controlled using a staircase to determine the interocular contrast ratio a participant requires to report seeing the letters in the left and right eye with equal probability.

We assessed balance point for three letter sizes: 1, 2, and 4 cpd; however, not all observers with amblyopia, particularly those with poor acuity, could conduct this task at the smaller letter sizes. Because of the high correlation among these measures (*r* ≥ 0.73) and the fact that there were no significant differences among them (*p* ≥ 0.31), we took the average balance point of as many measures as each participant was able to conduct. One participant with amblyopia was unable to conduct the task at any letter size (possibly due to amblyopic eye acuity of 1.06 logMAR). Data were not collected from 4 control participants.

### 4.6 Statistical Data analysis

The results of our visual assessment tasks (acuity, stereoacuity, contrast sensitivity and the interocular balance point) and model fits (k_AE,_ μ_AE,_ μ_FE,_ σ, and mean square error in fit) were analysed using t-tests (with Welch correction when appropriate) and ANOVA analyses using group and eye as factors. Effect sizes for pairwise tests are reported using Glass’s Δ using the control group’s standard deviation for reference. Because of the low sample number of strabismus with equal vision participants (n = 8), these data are described for comparison; formal statistics only compare performance between amblyopia and controls.

To determine whether the parameters of our three-stage model could provide predictive information about participants performance on visual assessment tasks we carried out hierarchical linear regression. In step 1 we predicted visual assessment measurements as a function of group membership alone. In step 2 we examined whether the addition of a model parameter as a predictor improved the regression predictions. We used a nested F-test to determine if the increase in R^2^ between steps 1 and 2 was statistically significant. All participants were used in this analysis, including strabismus with equal vision.

Statistical tests were carried out using R 4.0.4 (R Core Team, 2021), with the aid of the packages car^50^ (v 3.0-11), rstatix^51^ (v0.7.0), and tidyr^52^ (v1.1.3).

## 5 ACKNOWLEDGEMENTS

Research to Prevent Blindness Walt and Lilly Disney Award for Amblyopia Research (K.M./I.F.); Knights Templar Eye Foundation (K.M.); National Eye Institute and Office of Director, Office of Behavioral and Social Sciences Research R01EY014645 (I.F.); Natural Sciences and Engineering Council of Canada (K.M.); unrestricted grant from Research to Prevent Blindness to the University of Washington Department of Ophthalmology (K.T-H.).

## 7 SUPPLEMENTARY MATERIALS

### 7.1 Supplementary Tables

**Supplementary Table 1.**
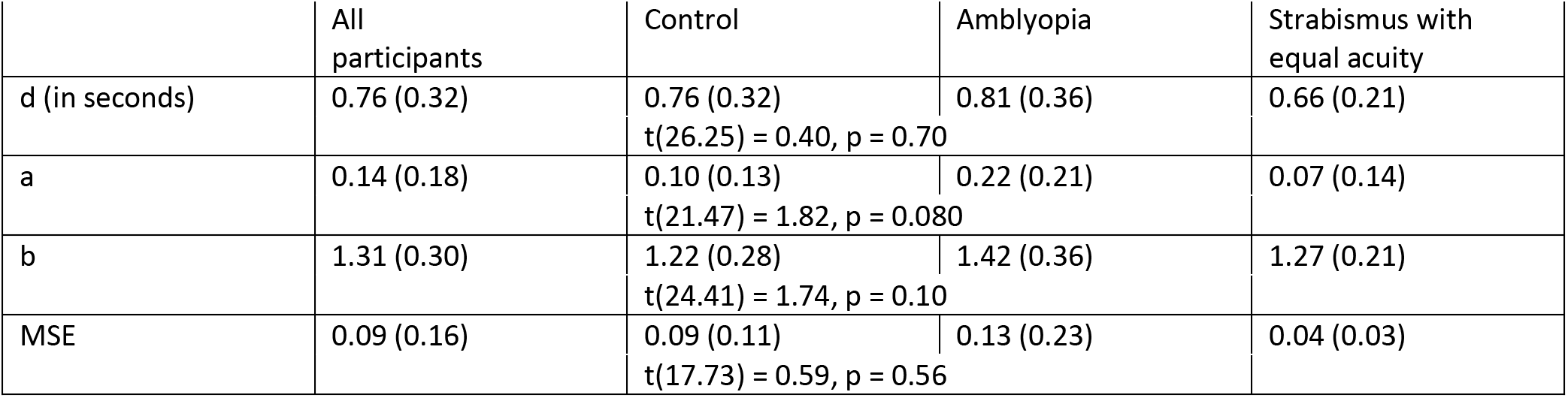
Stage 1 fits: Joystick calibration parameters. Mean (SD) values are shown. Statistical tests compare control vs. amblyopia groups.

|  | All participants | Control | Amblyopia | Strabismus with equal acuity |
| --- | --- | --- | --- | --- |
| d (in seconds) | 0.76 (0.32) | 0.76 (0.32)<br>$t(26.25) = 0.40, p = 0.70$ | 0.81 (0.36) | 0.66 (0.21) |
| a | 0.14 (0.18) | 0.10 (0.13)<br>$t(21.47) = 1.82, p = 0.080$ | 0.22 (0.21) | 0.07 (0.14) |
| b | 1.31 (0.30) | 1.22 (0.28)<br>$t(24.41) = 1.74, p = 0.10$ | 1.42 (0.36) | 1.27 (0.21) |
| MSE | 0.09 (0.16) | 0.09 (0.11)<br>$t(17.73) = 0.59, p = 0.56$ | 0.13 (0.23) | 0.04 (0.03) |

**Supplementary Table 2.** Linear regression results using Group and/or *k_AE_* as predictors.

| | Group alone | $k_{AE}$ alone | $k_{AE}$ after Group | Regression models (control group set as reference group) |
| --- | --- | --- | --- | --- |
| Acuity | $F(2,31) = 32.74, p < 0.001 *$ | $F(1,32) = 8.84, p = 0.0056 *$ | $F(1,30) = 0.25, p = 0.62$ | |
| Stereoacuity (log10) | $F(2,34) = 22.51, p < 0.001 *$ | $F(1,35) = 21.44, p < 0.001 *$ | $F(1,33) = 5.03, p = 0.032 *$ | $b_0 = 2.30$ (SE = 0.36), $t(33) = 6.47, p < 0.0001$; $b_{amblyopia} = 0.74$ (SE = 0.19), $t(33) = 3.91, p = 0.0004$; $b_{strabismus} = 0.80$ (SE = 0.19), $t(33) = 4.19, p = 0.0002$; $b_{k_{AE}} = -0.85$ (SE = 0.38), $t(33) = 2.24, p = 0.032$; $R^2 = 0.63, R^2_{adj} = 0.59, F(3,33) = 18.46, p < 0.0001$. |
| Contrast sensitivity | $F(2,29) = 10.73, p < 0.001 *$ | $F(1,30) = 5.70, p = 0.024 *$ | $F(1,28) = 0.99, p = 0.33$ | |
| Interocular balance point | $F(2,29) = 15.85, p < 0.001 *$ | $F(1,30) = 14.56, p < 0.001 *$ | $F(1,28) = 5.88, p = 0.022 *$ | $b_0 = 0.87$ (SE = 0.10), $t(28) = 8.52, p < 0.0001$; $b_{amblyopia} = 0.16$ (SE = 0.06), $t(28) = 2.69, p = 0.012$; $b_{strabismus} = -0.09$ (SE = 0.06), $t(28) = 1.58, p = 0.13$; $b_{k_{AE}} = -0.26$ (SE = 0.11), $t(28) = 2.42, p = 0.022$; $R^2 = 0.61, R^2_{adj} = 0.56, F(3,28) = 14.30, p < 0.0001$. |

Note. \* Statistically significant multiple regression model.

**Supplementary Table 3.** Linear regression results using μ_AE_ or μ_FE_ as predictors.

| | $\mu_{AE}$ alone | $\mu_{AE}$ after Group | $\mu_{FE}$ alone | $\mu_{FE}$ after Group |
| --- | --- | --- | --- | --- |
| Acuity (amblyopic eye) | $F(1,32) = 6.20, p = 0.018 *$ | $F(1,30) = 2.54, p = 0.12$ | $F(1,32) = 6.13, p = 0.019 *$ | $F(1,30) = 0.84, p = 0.37$ |
| Stereoacuity (log10) | $F(1,35) = 2.83, p = 0.10$ | $F(1,33) = 0.86, p = 0.36$ | $F(1,35) = 0.94, p = 0.34$ | $F(1,33) = 0.95, p = 0.34$ |
| Contrast sensitivity (amblyopic eye AUC) | $F(1,30) = 5.49, p = 0.026 *$ | $F(1,28) = 1.57, p = 0.22$ | $F(1,30) = 2.72, p = 0.11$ | $F(1,28) = 0.00, p = 0.95$ |
| Interocular balance point | $F(1,30) = 1.29, p = 0.27$ | $F(1,28) = 0.10, p = 0.76$ | $F(1,30) = 4.49, p = 0.042 *$ | $F(1,28) = 0.85, p = 0.37$ |

Note. \* Statistically significant multiple regression model.

**Supplementary Table 4.** Clinical details of participants with amblyopia and/or strabismus.

| ID | Subtype | Age | Acuity |  |  | Randot Circles (arcsec) | Treatment history | Refractive correction worn for testing |
| --- | --- | --- | --- | --- | --- | --- | --- | --- |
|  |  |  | BE | AE | FE |  |  |  |
| A1 | aniso | 25 | -0.06 | 0.23 (R) | -0.10 | 30 | Patching | R: +3.00 +1.00 x83<br>L: -3.00 |
| A2 § | aniso | 41 | -0.05 | 0.60 (L) | -0.12 | 140 | Patching | R: +3.25 +0.50 x10<br>L: +4.25 +0.75 x165 |
| A3 § | aniso | 46 | 0.06 | 0.56 (L) | 0.36 | 140 | No patching | R: -6<br>L: none ‡ |
| A4 | aniso | 71 | -0.11 | 0.23 (L) | -0.07 | 100 | Patching | R: +3.25 +1.5 x166<br>L: +5.25 +1.0 x135 |
| A5 * | aniso + strab | 18 | 0.11 | 0.33 (R) | 0.09 | 200 | Patching; No Sx | R: +5.50 -1.75 x170<br>L: +4.75 -1.25 x180 |
| A6 | aniso + strab | 23 | -0.21 | 1.06 (R) | -0.21 | Nil | Patching; No Sx | R: +1.0 +0.25 x27<br>L: -1.75 +0.25 x99 |
| A7 | aniso + strab | 42 | -0.11 | 0.55 (R) | -0.22 | Nil | Patching; 7 Sx | R: -5.5 +3.25 x82<br>L: -7.0 +2.50 x82 |
| A8 | aniso + strab | 61 | 0.02 | 0.18 (R) | -0.01 | Nil | Patching; 1 Sx | R: +1.50<br>L: +5.50 ‡ |
| A9 ¶ | aniso + strab | 70 | -0.11 | 0.79 (L) | -0.13 | Nil | Patching; 2 Sx | R: -1.5 +1.25 x145<br>L: plano |
| A10 | aniso + strab | 72 | 0.51 | 0.92 (R) | 0.67 | Nil | Patching; 3 Sx | R: plano<br>L: -2.50 +1.75 x155 |
| A11 | aniso + strab | 75 | 0 | 0.33 (L) | -0.03 | Nil | Patching; No Sx | R: -1.00 +2.00 x15<br>L: plano +1.25 x155 |
| A12 § | strab | 46 | 0.02 | 0.95 (R) | 0.04 | Nil | No patching; no Sx | R: +4.75 +0.50 x104<br>L: +4.00 |
| A13 § | strab | 51 | -0.04 | 0.96 (L) | -0.06 | Nil | Patching; 4 Sx | none |
| A14 | strab | 66 | 0.09 | 0.55 (L) | 0.19 | 140 | Patching; 2 Sx | R: -0.5 +0.5 x97<br>L: -0.75 +1.00 x103 |
| S1 | strab (equal vision) | 18 | 0.05 | 0.13 (L) | 0.02 | 100 | Patching; 3 Sx | R: -5.00 +2.25 x85<br>L: -3.75 +1.75 x85 |
| S2 § | strab (equal vision) | 22 | -0.08 | 0.00 (L) | -0.04 | Nil | Patching; 3 Sx | R: -6.5<br>L: -6.75 ‡ |
| S3 | strab (equal vision) | 22 | -0.02 | 0.08 (L) | 0.05 | Nil | Patching; 2 Sx | R: +4.00<br>L: +4.75 ‡ |
| S4 | strab (equal vision) | 36 | -0.19 | -0.08 (R) | -0.11 | 20 | No patching; no Sx | R: -2.25<br>L: -2.25 ‡ |
| S5 | strab (equal vision) | 50 | -0.16 | -0.14 (L) | -0.06 | Nil | Patching; 3 Sx | R: -1.75 +0.75 x95<br>L: -2.00 +0.25 x65 |
| S6 | strab (equal vision) | 50 | -0.03 | 0.01 (L) | -0.02 | 400 | No patching; no Sx | R: -8.75 +0.50 x170<br>L: -10.00 +1.50 x15 |
| S7 § | strab (equal vision) | 59 | N/A† | N/A† | N/A† | 200 | No patching; no Sx | R: -0.50<br>L: -0.25 ‡ |
| S8 | strab (equal vision) | 65 | 0.15 | 0.20 (L) | 0.22 | Nil | Patching; 2 Sx | R: -0.5 +0.75 x2<br>L: +0.5 +0.5 x7 |

**Note.** Aniso = anisometropia; strab = strabismus; BE = both eyes; AE = amblyopic eye; FE = fellow eye. The amblyopic eye of participants in the non-amblyopic strabismus with equal vision group reflects the eye assigned as amblyopic, as described in the main text. Unless noted otherwise, participants' residual undercorrection was <1 D spherical equivalent anisometropia and <1 D anisoastigmatism. \*Participant was assigned k=1 for both the amblyopic and fellow eyes (see text Section 4.3.2). †No acuity data available due to COVID-19 disruption. ‡Wearing contact lenses. §1.0 – 1.5 D uncorrected spherical equivalent anisometropia. ¶4.5 D undercorrected spherical equivalent anisometropia and 1.5 D uncorrected anisoastigmatism.

### 7.2 Supplementary Figures

**Supplementary Figure 1.**
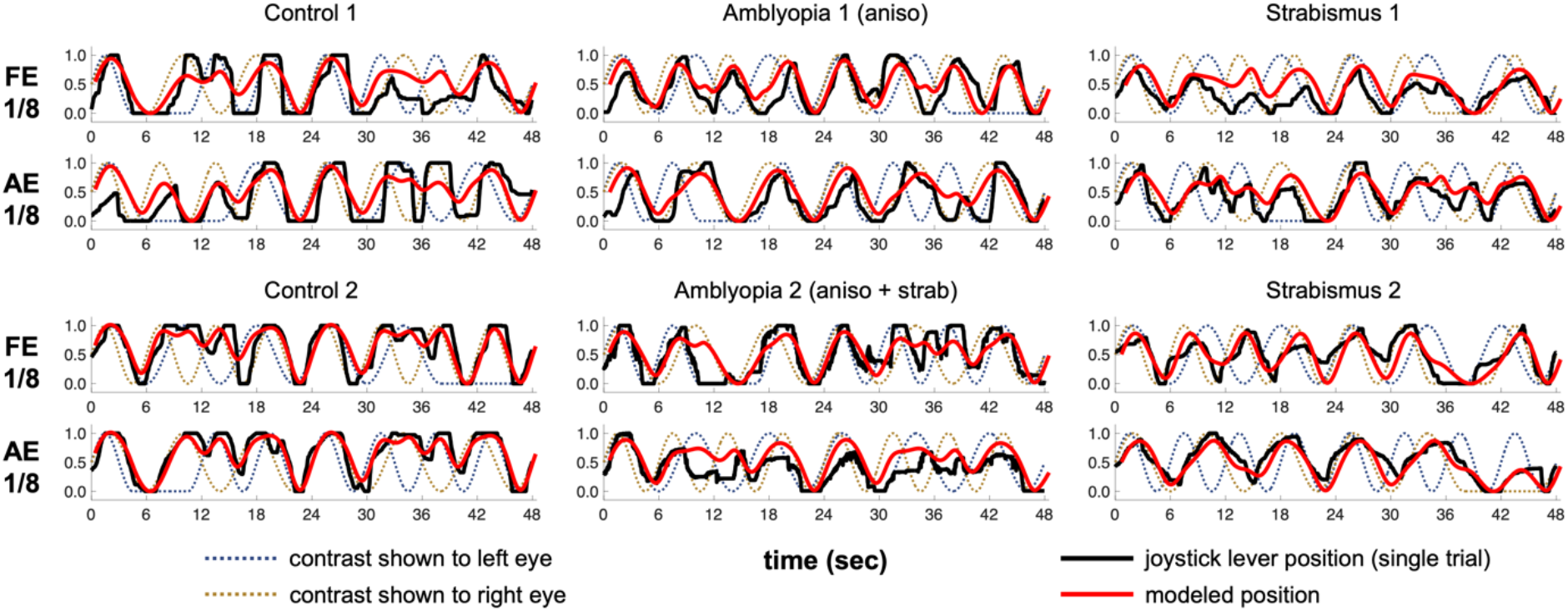
Example individual time-courses for single trials with full model fits, for typical participants from the control (left column), amblyopia (middle column) and non-amblyopic strabismus with equal vision (right column) groups. Participant 1 from each group corresponds to the participants shown in Figure 2C. For each participant, the top trace shows one trial where the participant’s “fellow” eye received 1/8 Hz contrast modulation and the “amblyopic” eye received 1/6 Hz; the bottom trace shows vice versa. The top and bottom observers with amblyopia correspond to A2 and A5, respectively, from Supplementary Table 4; the top and bottom observers with strabismus correspond to S5 and S8.

**Supplementary Figure 2.**
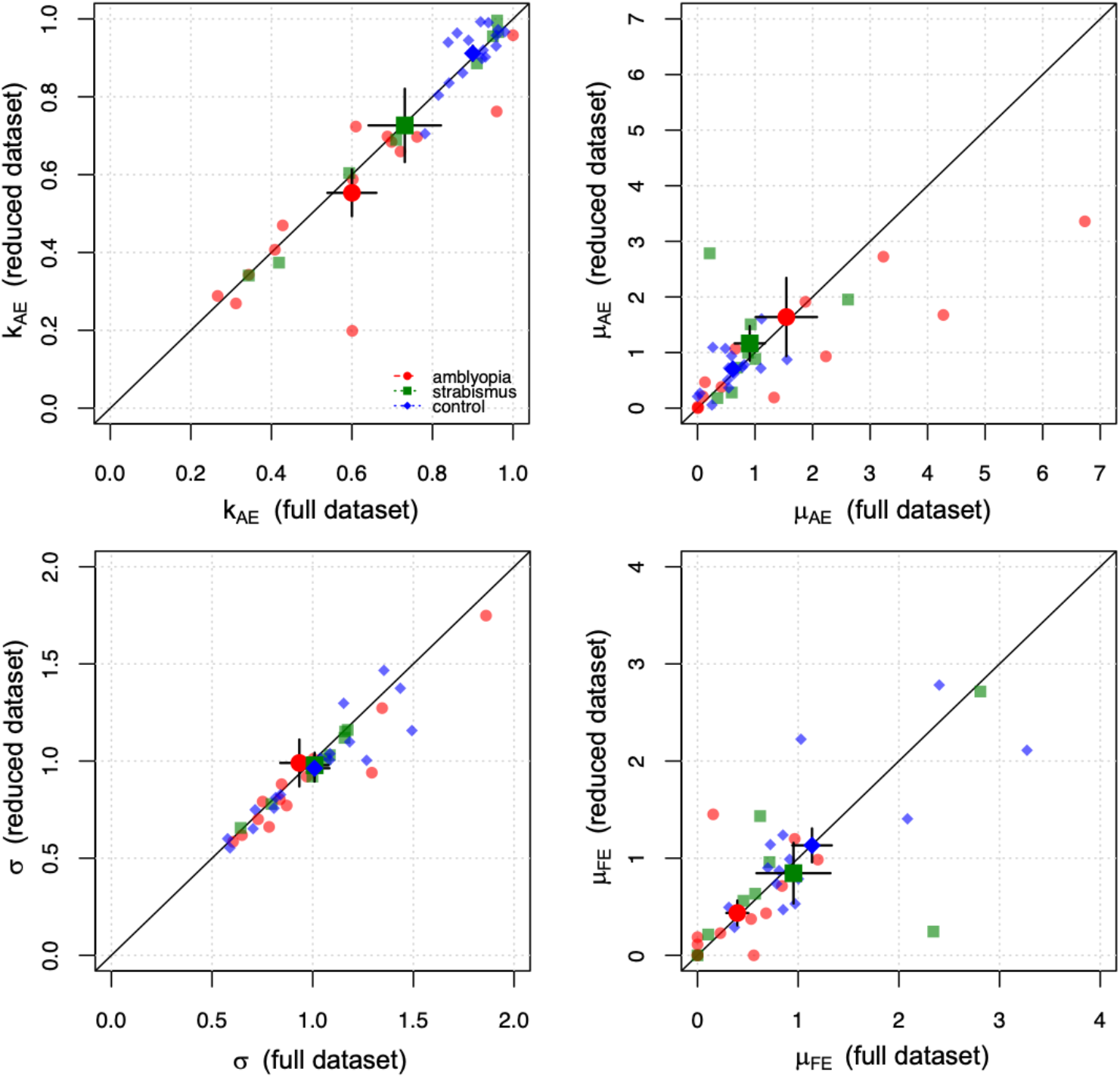
The relationship between the original estimates based on the full dataset (x-axis) and the estimates obtained from the reduced dataset (y-axis). Mean values and single standard errors are shown with the larger symbols (some error bars are too small to be seen). Individual data points are shown with the smaller symbols. The diagonal line represents unity.

### 7.3 Supplementary Data

Data and scripts for analysis are available at the UW’s Vision and Cognition Group github repository, which can be found at: https://github.com/VisCog

## Notes

### Competing Interest Statement

The authors have declared no competing interest.

## REFERENCES

1. Hu, B. et al. The global prevalence of amblyopia in children: A systematic review and meta-analysis. Front Pediatr 10, 819998 (2022).

2. Holmes, J. M. & Clarke, M. P. Amblyopia. Lancet 367, 1343–1351 (2006).

3. Kiorpes, L. & McKee, S. P. Neural mechanisms underlying amblyopia. Curr Opin Neurobiol 9, 480–486 (1999).

4. Birch, E. E. Amblyopia and binocular vision. Prog Retin Eye Res 33, 67–84 (2013).

5. Levi, D. M., Knill, D. C. & Bavelier, D. Stereopsis and amblyopia: A mini-review. Vision Res 114, 17–30 (2015).

6. Simons, K. Amblyopia characterization, treatment, and prophylaxis. Surv Ophthalmol 50, 123–166 (2005).

7. Baker, D. H., Meese, T. S. & Hess, R. F. Contrast masking in strabismic amblyopia: attenuation, noise, interocular suppression and binocular summation. Vision Res 48, 1625–1640 (2008).

8. Ding, J., Klein, S. A. & Levi, D. M. Binocular combination in abnormal binocular vision. J Vis 13, 14 (2013).

9. Ding, J. & Sperling, G. A gain-control theory of binocular combination. Proc Natl Acad Sci U S A 103, 1141–1146 (2006).

10. Meese, T. S., Georgeson, M. A. & Baker, D. H. Binocular contrast vision at and above threshold. J Vis 6, 1224–1243 (2006).

11. Moradi, F. & Heeger, D. J. Inter-ocular contrast normalization in human visual cortex. J Vis 9, 13.1–1322 (2009).

12. Bossi, M., Hamm, L. M., Dahlmann-Noor, A. & Dakin, S. C. A comparison of tests for quantifying sensory eye dominance. Vision Res 153, 60–69 (2018).

13. Huang, C. B., Zhou, J., Lu, Z. L. & Zhou, Y. Deficient binocular combination reveals mechanisms of anisometropic amblyopia: signal attenuation and interocular inhibition. J Vis 11, 4 (2011).

14. Zhou, J., Huang, P. C. & Hess, R. F. Interocular suppression in amblyopia for global orientation processing. J Vis 13, 19 (2013).

15. Lunghi, C., Morrone, M. C., Secci, J. & Caputo, R. Binocular rivalry measured 2 hours after occlusion therapy predicts the recovery rate of the amblyopic eye in anisometropic children. Invest Ophth Vis Sci 57, 1537–1546 (2016).

16. Mansouri, B., Thompson, B. & Hess, R. F. Measurement of suprathreshold binocular interactions in amblyopia. Vision Res 48, 2775–2784 (2008).

17. Kwon, M., Wiecek, E., Dakin, S. C. & Bex, P. J. Spatial-frequency dependent binocular imbalance in amblyopia. Sci Rep 5, 17181 (2015).

18. Katz, L. M., Levi, D. M. & Bedell, H. E. Central and peripheral contrast sensitivity in amblyopia with varying field size. Doc Ophthalmol 58, 351–373 (1984).

19. Dorr, M. et al. Binocular Summation and Suppression of Contrast Sensitivity in Strabismus, Fusion and Amblyopia. Front Hum Neurosci 13, 234 (2019).

20. Reynaud, A. & Hess, R. F. Is Suppression Just Normal Dichoptic Masking? Suprathreshold Considerations. Invest Ophthalmol Vis Sci 57, 5107–5115 (2016).

21. Birch, E. E. et al. Assessing Suppression in Amblyopic Children With a Dichoptic Eye Chart. Invest Ophthalmol Vis Sci 57, 5649–5654 (2016).

22. Beylerian, M. et al. Interocular suppressive interactions in amblyopia depend on spatial frequency. Vision Res 168, 18–28 (2020).

23. Zhou, J. et al. Amblyopic Suppression: Passive Attenuation, Enhanced Dichoptic Masking by the Fellow Eye or Reduced Dichoptic Masking by the Amblyopic Eye. Invest Ophthalmol Vis Sci 59, 4190–4197 (2018).

24. Gong, L. et al. Interocular Suppression as Revealed by Dichoptic Masking Is Orientation-Dependent and Imbalanced in Amblyopia. Invest Ophthalmol Vis Sci 61, 28 (2020).

25. Hess, R. F. & Bradley, A. Contrast perception above threshold is only minimally impaired in human amblyopia. Nature 287, 463–464 (1980).

26. Loshin, D. S. & Levi, D. M. Suprathreshold contrast perception in functional amblyopia. Doc Ophthalmol 55, 213–236 (1983).

27. Georgeson, M. A. & Sullivan, G. D. Contrast constancy: deblurring in human vision by spatial frequency channels. J Physiol 252, 627–656 (1975).

28. Peli, E. Suprathreshold contrast perception across differences in mean luminance: effects of stimulus size, dichoptic presentation, and length of adaptation. J Opt Soc Am A Opt Image Sci Vis 12, 817–823 (1995).

29. Webster, M. A., Werner, J. S. & Field, D. J. in Fitting the Mind to the World: Adaptation and After-Effects in High-Level Vision (eds Clifford, C. W. G. & Rhodes, G.) 241–278 (Oxford University Press, 2005).

30. Georgeson, M. A. Contrast overconstancy. J Opt Soc Am A 8, 579–586 (1991).

31. Legge, G. E. & Yuanchao, G. Stereopsis and contrast. Vision Res 29, 989–1004 (1989).

32. Webber, A. L., Schmid, K. L., Baldwin, A. S. & Hess, R. F. Suppression Rather Than Visual Acuity Loss Limits Stereoacuity in Amblyopia. Invest Ophthalmol Vis Sci 61, 50 (2020).

33. Shooner, C. et al. Asymmetric Dichoptic Masking in Visual Cortex of Amblyopic Macaque Monkeys. J Neurosci 37, 8734–8741 (2017).

34. Hess, R. F. Reasons why we might want to question the use of patching to treat amblyopia as well as the reliance on visual acuity as the primary outcome measure. BMJ Open Ophthalmol 7, e000914 (2022).

35. Mao, Y. et al. Binocular imbalance in amblyopia depends on spatial frequency in binocular combination. Invest Ophthalmol Vis Sci 61, 7 (2020).

36. Ding, J. & Levi, D. M. Binocular combination of luminance profiles. J Vis 17, 4 (2017).

37. Puckett, A. M. et al. Manipulating the structure of natural scenes using wavelets to study the functional architecture of perceptual hierarchies in the brain. NeuroImage 221, 117173 (2020).

38. Min, S. H. et al. A clinically convenient test to measure binocular balance across spatial frequency in amblyopia. iScience 25, 103652 (2022).

39. Martín, S., Portela, J. A., Ding, J., Ibarrondo, O. & Levi, D. M. Evaluation of a Virtual Reality implementation of a binocular imbalance test. PLoS One 15, e0238047 (2020).

40. Narasimhan, S., Harrison, E. R. & Giaschi, D. E. Quantitative measurement of interocular suppression in children with amblyopia. Vision Res 66, 1–10 (2012).

41. Chen, Y. et al. Patching and Suppression in Amblyopia: One Mechanism or Two. Front Neurosci 13, 1364 (2019).

42. Kelly, K. R. et al. Binocular iPad Game vs Patching for Treatment of Amblyopia in Children: A Randomized Clinical Trial. JAMA Ophthalmol 134, 1402–1408 (2016).

43. Yao, J., Moon, H. W. & Qu, X. Binocular game versus part-time patching for treatment of anisometropic amblyopia in Chinese children: a randomised clinical trial. Br J Ophthalmol 104, 369–375 (2020).

44. Brainard, D. H. The Psychophysics Toolbox. Spatial Vision 10, 433–436 (1997).

45. Pelli, D. G. The VideoToolbox software for visual psychophysics: transforming numbers into movies. Spatial Vision 10, 437–442 (1997).

46. Kleiner, M., Brainard, D. H. & Pelli, D. What’s new in Psychtoolbox-3? Perception 36, 14 (2007).

47. Bach, M. The Freiburg Visual Acuity Test-variability unchanged by post-hoc re-analysis. Graefes Arch Clin Exp Ophthalmol 245, 965–971 (2007).

48. Bach, M. The Freiburg Visual Acuity Test-automatic measurement of visual acuity. Optometry Vision Sci 73, 49–53 (1996).

49. Lesmes, L. A., Lu, Z. L., Baek, J. & Albright, T. D. Bayesian adaptive estimation of the contrast sensitivity function: the quick CSF method. J Vis 10, 17.1–1721 (2010).

50. Fox, J. & Weisberg, S. An R companion to applied regression (Sage, Thousand Oaks, CA, 2019).

51. Kassambara, A. rstatix: Pipe-friendly framework for basic statistical tests. R package version 0.7.0, (2021).

52. Wickham, H. tidyr: Tidy Messy Data. R package version 1.1.3, (2021).

